# Prenatal exposure to THC vapor influences feeding, bodyweight and glucose metabolism under both basal conditions and following high fat diet

**DOI:** 10.1101/2023.09.29.560189

**Authors:** Catherine Hume, Samantha L. Baglot, Lucia Javorcikova, Savannah H.M. Lightfoot, Jessica B.K. Scheufen, Matthew N. Hill

## Abstract

4-20% of people report using cannabis during pregnancy, thereby it is essential to assess the associated risks. There is some evidence that prenatal cannabis exposure (PCE) may be associated with increased risk for development of obesity and diabetes later in life, however this has not been well explored under controlled conditions. The aim of this study was to use a translational THC vapor model in rodents to characterize the effects of PCE on adiposity, glucose metabolism, and feeding patterns in adulthood, with focus on potential sex differences. Pregnant Sprague Dawley rats were exposed to vaporized THC (100mg/ml) or control (polyethylene glycol vehicle) across the entire gestational period. Adult offspring from PCE or control litters were subjected to measures of adiposity, glucose metabolism and feeding behavior. Rats were then placed onto special diets (60% high-fat diet [HFD] or control 10% low fat diet [LFD]) for 4-months, then re-subjected to adiposity, glucose metabolism and feeding behavior measurements. PCE did not influence maternal weight or food consumption but was associated with transient decreased pup weight. PCE did not initially influence bodyweight or adiposity, but PCE did significantly reduce the rate of bodyweight gain when animals were maintained on special diets (HFD/LFD), regardless of which diet. Further, PCE had complex effects on glucose metabolism and feeding behavior that were both sex and diet dependent. No effects of PCE were found on plasma leptin or insulin, or white adipose tissue mass. Overall, this data enhances current understanding of the potential impacts of PCE.

## Introduction

Around 20% of people, 16 years or older, in North America report using cannabis in the last month (1). With legalization of cannabis in Canada and parts of the United States, the availability, demand, and acceptance surrounding cannabis use has increased. In fact, 70% of people of reproductive age believe that there is little or no harm in using cannabis on a weekly basis (2). It is reported that between 4-10% of people use cannabis during pregnancy, with those under the age of 24 reporting usage as high as 22% (2, 3); further, as positive toxicology levels are even higher than self-report, these number are likely an underrepresentation (3). Cannabis is utilized during pregnancy for managing pregnancy-related side effects (such as nausea, pain, and sleep), for managing pre-existing conditions (such as anxiety), and for recreational purposes (4, 5). Many pregnant people believe cannabis is safer to use than prescription or over-the-counter medications and a large proportion of dispensaries recommend using cannabis products to manage morning sickness (6), despite a considerable lack of knowledge surrounding the impacts of gestational exposure to cannabinoids.

With increased perceptions of safety, elevated potency of delta-9-tetrahydrocannabinol (THC; the main psychoactive component of cannabis) (7, 8), and the ability for THC and its metabolites to cross the placenta (9, 10), the effects of prenatal cannabis exposure (PCE) on developing offspring are important considerations. Human and rodent studies of PCE have found substantial evidence for lower birth weight (11–21), as well as some evidence for executive function, attention, and cognition changes (13, 22-26); increased incidence of autism (22) or reduced social interaction (21, 27); earlier age and increased likelihood of substance use (13, 28-30) or altered drug-seeking behaviour (31); and finally, dysregulated emotional behaviour (13, 32, 21, 27). However, many of these studies have reported opposing results and/or little to no effects on many other domains, and therefore should be interpreted with caution. Human studies rely on self-report data, are confounded by an unknown timing and amount of cannabis exposure, and cannot eliminate the possible effects of exposure to stress and other drugs during pregnancy or adverse early-life environments. While animal models can control for these confounds, many models utilize injection administration of THC across a wide range of doses, some of which likely exceed typical exposure levels seen in human use (9, 33–37).

To date the most consistent PCE-related finding is a reduction in birth weight. Interestingly, a reduction in birth weight is common after other prenatal insults, such as prenatal alcohol or stress exposure, and is often associated with increased (or rebound) bodyweight and/or metabolic changes later in life (38). A few human studies following PCE have shown higher fat mass, adiposity, and fasting glucose in PCE children at ∼5 years old (14) and higher BMI at ∼10 years old (15, 39). These results provide some evidence for disrupted glucose homeostasis in childhood following PCE and suggest increased risk for the development of obesity and diabetes later in life (40), however no human study to date has looked beyond childhood. Interestingly, recent animal studies utilizing injections of THC examined the metabolic outcomes of PCE into adulthood and showed increased visceral adiposity (41), a reduction in fasting glucose but not fasting insulin in both males and females, and increased levels of glucose and insulin following dextrose injection (i.e. the glucose tolerance test) in females only (19). Another study found increased food motivation and reduced sucrose preference in males following the combination of PCE and acute stress exposure (42). Taken together, these studies suggest altered glucose homeostasis and metabolism, as well as altered feeding behaviour, following PCE.

This study sought to determine whether a translationally relevant *vapor model of PCE* increases male and female offspring’s risk for obesity, diabetes, and abnormal feeding habits. Further, our study is the first to examine whether PCE interacts with acute stress and high-fat diet (HFD) to influence energy balance.

## Methods

### Animals, Housing and Breeding

Adult male (n=12) and female (n=30) Sprague-Dawley rats were obtained from Charles River Laboratories (New York, United States) and pair-housed according to sex in standard caging (constant temperature [21 ± 1℃] on a 12-hr light-dark cycle [lights on at 07:00] with *ad libitum* standard chow and water). Breeding occurred in house with vaginal lavage performed each morning. The presence of sperm indicated gestation day 1 (G1) and body weight and food intake were measured weekly (G1, 7, 14, 21). All experiments were performed in accordance with the Canadian Council on Animal Care (CCAC) guidelines, and with the approval of the University of Calgary Animal Care Committee (ACC).

### Prenatal Cannabis Exposure

For 3 days prior to, and throughout breeding, females were habituated to the vapor chambers (15-min with only air flow) to reduce stress. On G1, females were single housed and randomly assigned to one of two treatment groups: control or THC-exposure. Control dams (n=14) were exposed to polyethylene glycol (PEG-400) vapor, whereas THC dams (n=14) were exposed to THC-dominant cannabis extract containing 89.7% THC, 0.3% CBD, 1.7% CBG, 1.4% CBN and 0.6% CBC (Aphria Inc., Ontario, Canada), diluted to 100 mg/ml THC in PEG-400. All dams were exposed via a validated (9, 33, 36) vapor chamber system (La Jolla Alcohol Research Inc., California, USA). Dams were exposed within their home cages, which fit directly into one of four compartments within the chamber, once per day between 09:00 and 12:00 hr until the day prior to expected delivery (G21). Compartment location was randomized each day. Exposure consisted of a 15-min session (10-sec puff once every 45-sec). To determine plasma THC levels, blood samples were taken from the tail vein of all dams on G1 and G10 immediately following exposure and underwent mass spectrometry analysis (9). Plasma THC levels for this breeding averaged at 109.8 ± 24.7 ng/ml on G1 and 182.0 ± 31.6 ng/ml on G10, indicating a moderate level of exposure.

### Birth, Culling and Weaning

The day after delivery (deemed postnatal day [P]1), dams and their litters were weighed, and pups were culled to 12 (6 females and 6 males whenever possible). If needed, pups born on the same day and from the same treatment group (control or THC-exposed) were cross fostered to maintain litter size and sex composition. Dam and litter weight were taken weekly (P1, 7, 14, 21). Pups were weaned on P22, split into same sex groups for ∼5 days, and then pair-housed on ∼P28. Individual bodyweights were taken at P50, P59, P68 and P77. Four weeks prior to testing offspring were transferred to a reverse 12-hr dark-light cycle room (lights off at 08:00) with *ad libitum* access to standard rodent chow (Prolab® RMH 2500; 3.34 kcal/g) and water. Two weeks prior to testing offspring were singly housed.

### General Experimental Design

Adult offspring were subjected to glucose metabolism and feeding behaviour tests between P50-P77 (referred to as “Chow Testing”), were then put on a high-fat diet (HFD) or low-fat control diet (LFD) for four-months between P78-P191 (referred to as “HFD Access”), and finally re-subjected to glucose metabolism and feeding behaviour tests between P192-220 (referred to as “HFD Testing”). Overall, 48 age-matched rats were used: 24 males (n=12 control and n=12 PCE) and 24 females (n=12 control and n=12 PCE). Half of these animals were placed on HFD and half on LFD. Bodyweight and energy intake was tracked across the experiment.

### Glucose Metabolism Testing

#### Intraperitoneal Glucose Tolerance Test (IPGTT)

Rats were fasted for 16-hrs. At timepoint (T) 0, blood glucose was measured via tail nick (Accu-Chek® Guide Meter, Roche) and rats were then injected with dextrose (2g/kg in saline, IP). Blood glucose was also measured at T15, 30, 60 and 120. Rats were habituated to handling and restraint procedures for three days prior to IPGTT.

#### Plasma Insulin Measurements

Plasma insulin levels were measured in basal and fasting conditions, as well as following dextrose injection. Blood was collected prior to the 16-hr fast (basal), at T0 (fasting), and at T30 (post dextrose injection). All blood was collected into EDTA coated tubes and stored on ice until centrifuged at 10,000 rpm for 20 min at 4°C, then plasma stored at −80°C until analysis. Plasma was processed using an insulin ELISA kit (Millipore Sigma; #EZRMI-13K) following manufacturer’s instructions in duplicate. T0 samples were ran undiluted, T30 and basal were diluted 1:1 in assay buffer. Intra-assay variability was <10%.

#### Homeostatic Model Assessment for Insulin Resistance (HOMA-IR)

HOMA-IR values were calculated on T0 (fasting) levels: T0 glucose (mmol/L) x T0 insulin (ng/mL)/22.5 (19).

### Feeding Behaviour Testing

#### Daily Energy Intake Pattern Measurements

Food was measured at the end of the light-phase (08:00) and every two hours throughout the dark-phase (10:00-20:00). Measurements were carried out for 3-days for habituation, and final data collected in one ‘test’ day.

#### Macronutrient-Specific Food Choice Test

During the dark-phase, rats were given access to both a high-fat food (80% fat, 10% carbohydrate, 10% protein; 6.1 kcal/g; #D19102309, Research Diets Inc., New Jersey, USA) and a high-carbohydrate food (80% carbohydrate, 10% fat, 10% protein; 3.8 kcal/g; #D19102310, Research Diets Inc.), and intake of each food was measured every two hours (10:00-20:00). A modified food hopper was used to ensure equal access to both foods, and the position of each food was switched daily to account for potential side preferences. Following 12-hour food choice rats were returned to their maintenance diet (chow or HFD/LFD dependent on phase of experiment) during the light-phase and intake measured. Measurements were carried out for 3-days to habituate animals, and final data collected in two ‘test’ days (43, 44).

#### Macronutrient-Specific Food Choice Following Stress

During the ‘Chow-Testing’ phase of the experiment, rats were subjected to a third food-choice ‘test’ day following 30-min restraint stress to examine the combinatorial effects of PCE and acute stress. Stress occurred directly prior to dark-phase onset (07:30-08:00) and food choice measurements occurred as described above.

#### Sucrose Preference Test

During the ‘HFD Testing’ phase of the experiment, a two-bottle choice test was used to assess sucrose preference (45). Rats were given *ad libitum* access to both a bottle of water and a bottle of 2% sucrose for 4-day for habituation. On test day, intake of each was measured during the first 2 hours of the dark-phase (08:00-10:00). Bottle position was randomized between animals to account for potential side preferences. Rats had food access throughout.

### High-fat Diet (HFD) Access

Following the ‘Chow-Testing’ phase, rats were given *ad libitum* access to a HFD (60% fat; 5.24 kcal/g; #D12492, Research Diets Inc., New Jersey, USA) or control LFD (10% fat, matching sucrose to #D12492; 3.85 kcal/g; #D12450J, Research Diets Inc.) for four months. Energy intake and bodyweight was measured every 3-days.

### Adiposity Assessment

#### Basal Plasma Leptin Measurements

To indirectly measure adiposity, basal plasma (collection prior to IPGTT) leptin was measured using a leptin ELISA (Millipore Sigma; #EZRL-83K) following manufacturer’s instructions in duplicate. ‘Chow-Testing’ basal plasma was run undiluted and ‘HFD Testing’ basal plasma was diluted 1:1 in assay buffer. Intra-assay variability was <10%.

#### White Adipose Tissue (WAT) Mass

To directly measure adiposity, WAT pad mass was measured at the end of the study. Rats were euthanized and subcutaneous inguinal, visceral retroperitoneal and gonadal fat pads were dissected bilaterally and weighed (46). Adipose to bodyweight ratio was calculated: total WAT mass (g) / bodyweight (g) (41).

### Statistics

All data is presented as mean ± SEM. Intake data is normalised to bodyweight and thereby shown as kcal/kg. One female PCE rat was removed from the entire dataset due to illness, and one male HFD-fed control rat was removed from the food choice and sucrose preference analyses due to a failure to eat high-carbohydrate food or drink sucrose. Data was analyzed using either a univariate ANOVA or a repeated-measures ANOVA. Repeated-measures ANOVAs consisted of either day, time, or stress as the within-subjects factor. For the chow-testing phase of the experiment, all analyses consisted of prenatal group (CON versus PCE) and sex (Male versus Female) as between-subjects factors. For the HFD testing phase of the experiment, all analyses consisted of prenatal group, sex, and diet (HFD versus LFD) as between-subjects factors. When necessary for clarity, male and female data is graphed separately despite being analyzed together. Post-hoc analyses were corrected for multiple comparisons using Bonferroni and repeated measures analyses were normalized using Greenhouse-Geisser if data violated Mauchly’s Test of Sphericity. Data was considered significant at p≤0.05.

## Results

### Maternal & Litter Outcomes

All dams gained weight across gestation (main effect of time, F_(2,184,56.787)_=949.89 at p<0.001; Table 1A), and all dams had greater energy intake between G7-G14 compared to G14-G21 (main effect of week, F_(2,40)_=8.727 at p<0.001; Table 1A). No prenatal group differences were found. No significant differences were found in litter size or litter weight at birth (data not shown).

**Table 1:** Maternal and Pup Outcomes and Basal Circulating Factors. **(A)** *Maternal outcomes*: maternal weight (top row; g) at G1, G7, G14 and G21 (main effect of time, F_(2.184,56.787)_=949.89 at p<0.001, #: all timepoints significantly different than all other timepoints) and weekly maternal food consumption (bottom row) for G1-G7, G7-G14 and G14-G21 (kcal/kg/day; main effect of week, F_(2,40)_=8.727 at p<0.001; #: G7-G14 > G14-G21). (B) *Pup weights:* male (top row) and female (bottom row) pup weights at P1, P8, P15 and P22 (g/#; interaction effect of age and prenatal group, F_(1.341,69.718)_=4.665 at p=0.024, *: PCE < CON at P22 at p=0.039; main effect of sex, F_(1,52)_=5.347 at p=0.25, $: all ages different to all other ages at p<0.001). **(C)** *Chow-fed basal circulating factors:* blood glucose (top row; main effect of sex, F_(1,47)_ = 29.539 at p<0.001, $: male > female), plasma insulin (middle row; main effect of sex, F_(1,47)_ = 56.211 at p<0.001, $: male > female) and plasma leptin (bottom row; main effect of sex, F_(1,47)_ = 17.865 at p<0.001, $: male > female) concentrations in male and female, PCE and control animals. **(D)** *HFD/LFD-fed basal circulating factors:* blood glucose (top row; interaction effect of prenatal group and sex, F_(1,,47)_= 4.282 at p=0.045, *: male PCE > male CON, ^: HFD < LFD, $: male > female), plasma insulin (middle row; interaction effect of prenatal group and diet, F_(1,47)_=4.060 at p=0.05, ^: CON HFD < CON LFD, $: male > female) and plasma leptin (bottom row; main effect of diet, F_(1,47)_ = 6.463 at p=0.015, ^: HFD > LFD; main effect of sex, F_(1,47)_ = 25.140 at p<0.001, $: male > female) concentrations in male and female, PCE and control animals.

| A. Maternal Outcomes |  |  |  |  |  |  |  |  |
| --- | --- | --- | --- | --- | --- | --- | --- | --- |
| Variable | G1 |  | G7 |  | G14 |  | G21 |  |
|  | CON | PCE | CON | PCE | CON | PCE | CON | PCE |
| Maternal Weight (g) <sup>#</sup> | 259.6 ± 3.3 | 257.9 ± 3.4 | 285.2 ± 4.0 | 277.2 ± 4.1 | 309.6 ± 5.6 | 309.2 ± 5.8 | 382.1 ± 6.4 | 382.8 ± 6.7 |
| Weekly Food Consumption (kcal/kg/day) | - | - | 201.3 ± 10.2 | 218.2 ± 10.2 | 239.7 ± 9.5<sup>#</sup> | 221.5 ± 9.5<sup>#</sup> | 190.6 ± 7.9 | 201.5 ± 7.9 |
| B. Pup Weights |  |  |  |  |  |  |  |  |
| Variable | P1<sup>#</sup> |  | P8<sup>#</sup> |  | P15<sup>#</sup> |  | P22<sup>#</sup> |  |
|  | CON | PCE | CON | PCE | CON | PCE | CON* | PCE* |
| Male<sup>\$</sup> Pup Weights (g/#) | 7.2 ± 0.1 | 7.3 ± 0.1 | 18.2 ± 0.4 | 18.0 ± 0.4 | 33.3 ± 0.5 | 32.3 ± 0.5 | 59.8 ± 1.2 | 57.0 ± 1.2 |
| Female<sup>\$</sup> Pup Weights (g/#) | 6.7 ± 0.1 | 6.9 ± 0.1 | 17.4 ± 0.4 | 17.4 ± 0.4 | 32.0 ± 0.5 | 31.5 ± 0.5 | 57.2 ± 1.2 | 54.7 ± 1.2 |
| C. Chow-Fed Basal Circulating Factors |  |  |  |  |  |  |  |  |
| Plasma Factor | Male |  | Female |  |  |  |  |  |
|  | CON | PCE | CON | PCE |  |  |  |  |
| Glucose (mmol/L) <sup>\$</sup> | 6.7 ± 0.1 | 6.8 ± 0.1 | 6.2 ± 0.1 | 6.1 ± 0.1 | | | | |
| Insulin (ng/mL) <sup>\$</sup> | 11.2 ± 0.8 | 10.1 ± 0.8 | 4.9 ± 0.8 | 5.1 ± 0.6 | | | | |
| Leptin (ng/mL) <sup>\$</sup> | 10.5 ± 0.5 | 10.7 ± 0.9 | 7.4 ± 0.8 | 7.7 ± 0.6 | | | | |
| D. HFD/LFD-Fed Basal Circulating Factors |  |  |  |  |  |  |  |  |
| Plasma Factor | LFD |  |  |  | HFD |  |  |  |
|  | Male |  | Female |  | Male |  | Female |  |
|  | CON | PCE | CON | PCE | CON | PCE | CON | PCE |
| Glucose (mmol/L) <sup>\$^</sup> | 6.5 ± 0.2 * | 6.8 ± 0.2 * | 6.1 ± 0.2 | 5.8 ± 0.2 | 6.1 ± 0.2 * | 6.5 ± 0.2 * | 5.7 ± 0.2 | 5.7 ± 0.2 |
| Insulin (ng/mL) <sup>\$</sup> | 23.2 ± 1.7 ^ | 23.0 ± 1.7 | 10.5 ± 1.7 ^ | 7.0 ± 1.7 | 13.9 ± 1.7 ^ | 18.4 ± 1.7 | 7.1 ± 1.7 ^ | 8.8 ± 1.9 |
| Leptin (ng/mL) <sup>\$^</sup> | 58.9 ± 7.4 | 53.9 ± 7.4 | 33.8 ± 7.4 | 24.0 ± 7.4 | 62.4 ± 7.4 | 75.2 ± 7.4 | 40.4 ± 7.4 | 46.3 ± 8.1 |

### Pup Bodyweight

PCE pups weighed less than control pups on P22 (p=0.039; interaction effect of age and prenatal group, F_(1.341,69.718)_=4.665 at p=0.024; Table 1B) but no significant differences were found for any other ages. Both PCE and control pups gained weight across the postnatal period (all ages differed at p<0.001, Table 1B), and males weighed more than females (main effect of sex, F_(1,52)_=5.347 at p=0.25; Table 1B).

### Adult Offspring Bodyweight

No significant differences in bodyweight (data not shown) or bodyweight gain (Figure 1A) were found across prenatal groups. Over time, all rats gained weight (p<0.001) and males gained more than females (p<0.001; interaction effect of age and sex, F_(1.25, 53.77)_ = 34.103 at p<0.001; Figure 1A).

**Figure 1:**
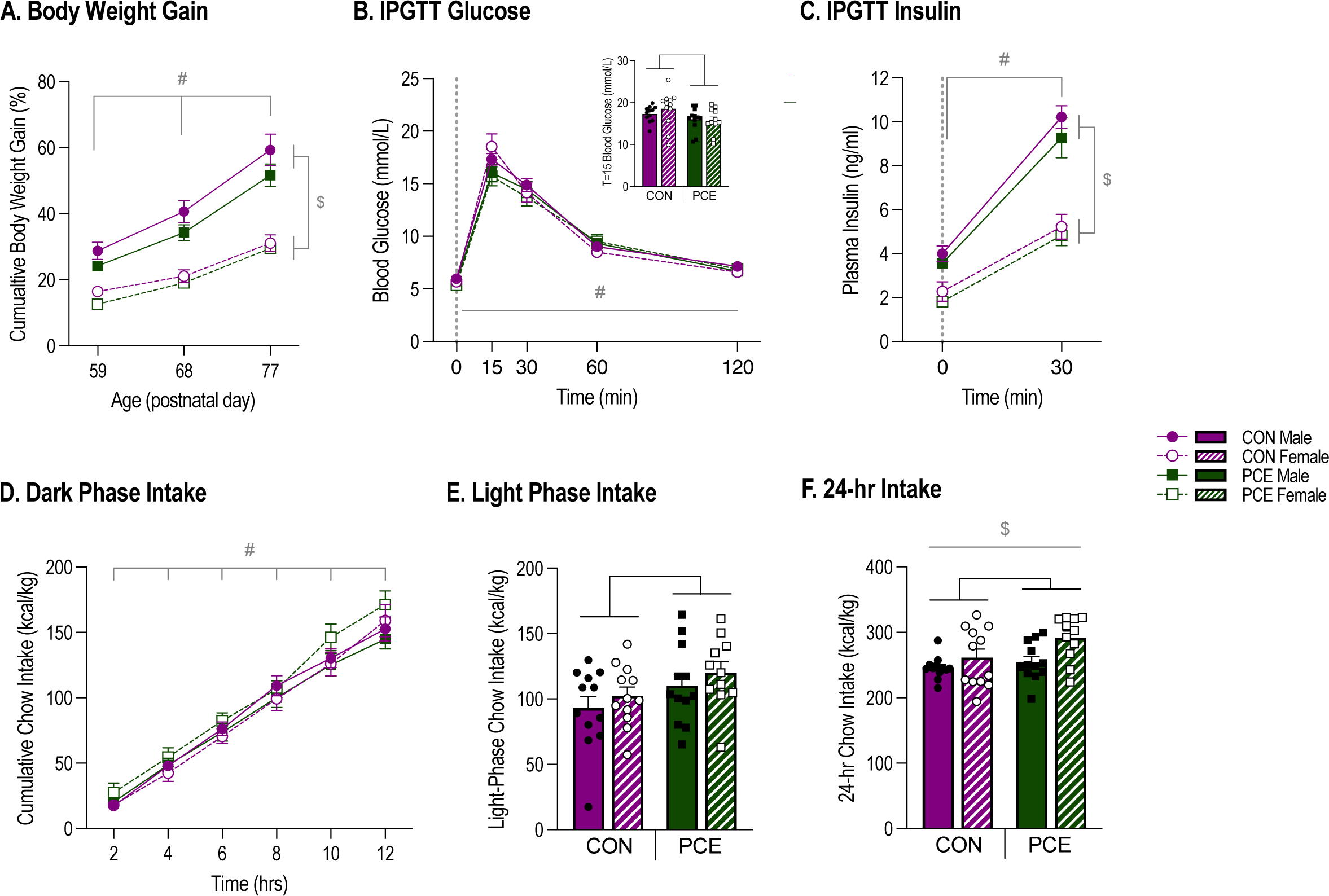
In chow-fed offspring, PCE has no effect on bodyweight gain, but improves glucose tolerance and increases 24-hr chow intake. (A) Cumulative bodyweight gain (%) from P50 (interaction of age and sex, F_(1.25, 53.77)_ = 34.103 at p<0.001, #: all ages significantly different than all other ages at p<0.001, $: males > females at every age at p<0.001). (B) Blood glucose concentrations (mmol/L) across the IPGTT; dextrose injection at T0 (interaction of time and prenatal group, F_(1.769, 76.046)_ = 26.693 at p=0.021, *: PCE < CON only at T15 at p=0.026, #: all timepoints significantly different than all other timepoints at p<0.001). Dotted line indicates timing of dextrose injection. *Inset:* T15 IPGTT blood glucose (mmol/L; *: PCE < CON at p=0.026). **(C)** Plasma insulin (ng/ml) at IPGTT T0 and T30; dextrose injection at T0 (interaction of time and sex, F_(1,43)_ = 19.294 at p<0.001, #: T30 > T0 at p<0.001, $: males > females at each time point at p<0.001). Dotted line indicates timing of dextrose injection. **(D)** Cumulative dark-phase chow intake (kcal/kg) over time (main effect of time, F_(2.727,117.261)_ = 573.242 at p<0.001, #: all timepoints significantly different than all other timepoints). **(E)** Total 12-hour light-phase chow intake (kcal/kg; main effect of prenatal group, F_(1,47)_ = 4.49 at p=0.04, *: PCE > CON). **(F)** 24-hr chow intake (kcal/kg; main effect of prenatal group, F_(1,47)_ = 4.237 at p=0.046, *: PCE > CON; main effect of sex, F_(1,47)_ = 7.465 at p=0.009, $: males < females).

### Chow-Testing: Basal Circulating Factors

For basal blood glucose, plasma insulin and plasma leptin concentrations, no significant differences were found between prenatal groups (Table 1C). Males showed higher levels of all three circulating factors than females (glucose: main effect of sex, F_(1,47)_ = 29.539 at p<0.001; insulin: main effect of sex, F_(1,47)_ = 56.211 at p<0.001; leptin: main effect of sex, F_(1,47)_ = 17.865 at p<0.001; Table 1C).

### Chow-Testing: Glucose Tolerance

For blood glucose, peak concentrations at T15 following dextrose injection were lower for PCE animals than controls (p=0.026; interaction of time and prenatal group, F_(1.769, 76.046)_=26.693 at p=0.021; Figure 1B). All other timepoints did not differ between prenatal groups (Figure 1B). Following dextrose injection, blood glucose for all animals followed the expected IPGTT time course (all timepoints different than all other timepoints at p<0.001; Figure 1B).

For IPGTT plasma insulin, no effects of prenatal group were found. As expected, insulin concentrations were higher after dextrose injection (p<0.001) and were higher in males than females (p<0.001; interaction of time and sex, F_(1,43)_=19.294 at p<0.001; Figure 1C).

For HOMA-IR, no effects of prenatal group were found. Males had higher HOMA-IR values than females (main effect of sex, F_(1,43)_=26.799 at p<0.001) (data not shown).

### Chow-Testing: Feeding Patterns

Dark-phase chow intake patterns were unaffected by prenatal group or sex, where all rats steadily consumed chow throughout the whole dark-phase (main effect of time, F_(2.727,117.261)_=573.242 at p<0.001; Figure 1D). Light-phase and 24-hr chow intake was higher for PCE animals than controls (light-phase: main effect of prenatal group, F_(1,47)_=4.49 at p=0.04; Figure 1E; 24-hr: main effect of prenatal group, F_(1,47)_=4.237 at p=0.046; Figure 1F). 24-hr chow intake was higher in females than males (main effect of sex, F_(1,47)_=7.465 at p<0.001; Figure 1F).

### Chow-Testing: Food Choice

At the 2-hour food choice timepoint, PCE animals consumed more high-carbohydrate food than control animals in both no stress and stress conditions (main effect of prenatal group, F_(1,43)_=4.123 at p=0.049; Figure 2A & Supplementary Figure 1A). No effects of prenatal group were found for high-fat food intake or total combined food intake (high-carbohydrate + high-fat food intake; Figure 2A & Supplementary Figure 1A).

**Figure 2:**
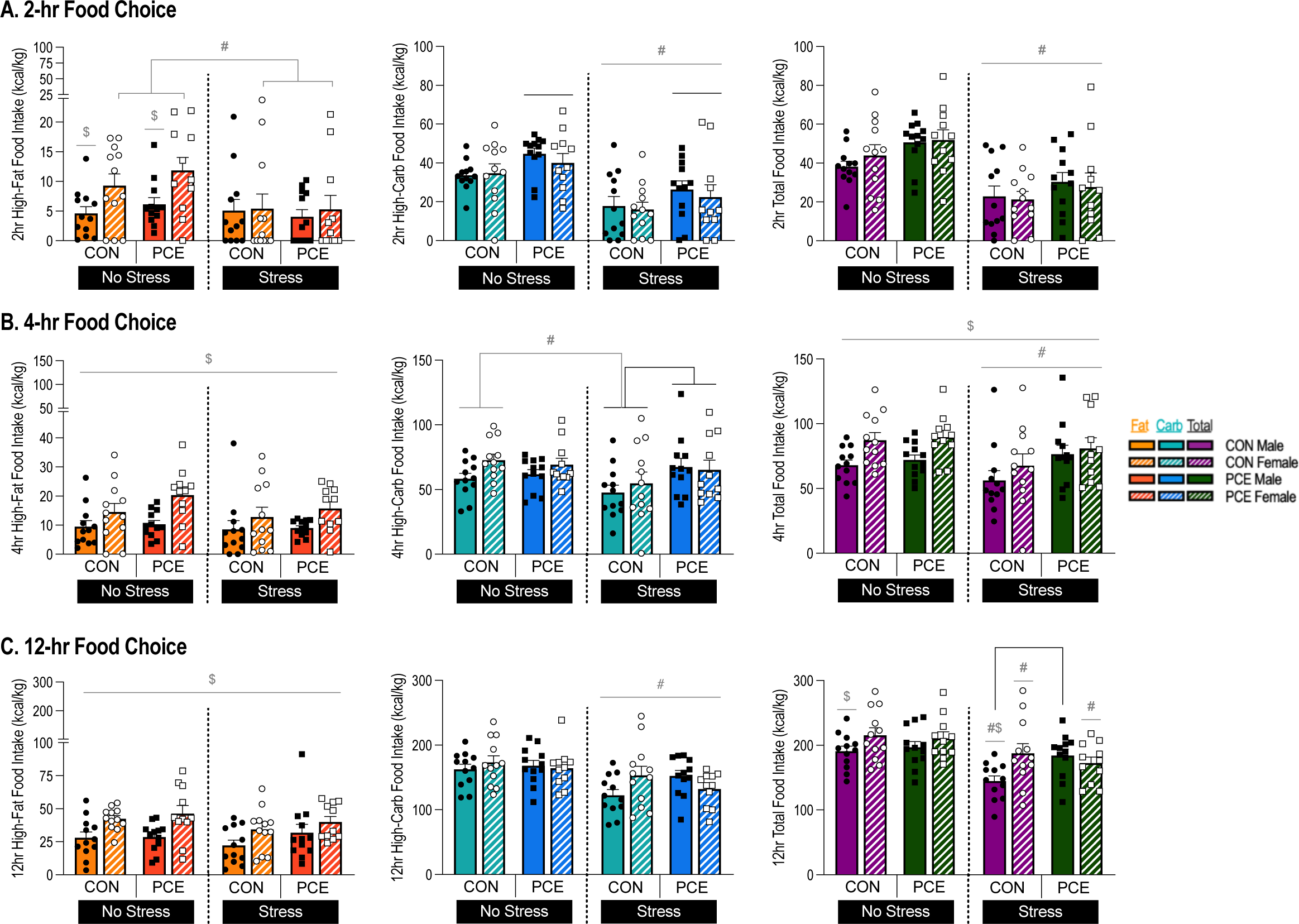
In chow-fed offspring, PCE acutely increases high-carbohydrate food intake when given a food choice, an effect that is prolonged by stress. **(A)** 2-hour food choice timepoint intake (kcal/kg) of high-fat food (left panel; interaction effect of sex and stress, F_(1,43)_ = 4.534 at p=0.039, #: female no stress > female stress at p=0.001, $: male < female at p=0.003), high-carbohydrate food (center panel; main effect of prenatal group, F_(1,43)_ =4.123 at p=0.049, *: PCE > CON; main effect of stress, F_(1,43)_ = 72.820 at p<0.001, #: no stress > stress), and total combined food (high-fat + high-carbohydrate food; right panel; main effect of stress, F_(1,43)_ = 91.390 at p<0.001, #: no stress > stress) during the food choice test, both in no stress and stress conditions. **(B)** 4-hour food choice timepoint intake (kcal/kg) of high-fat food (left panel; main effect of sex, F_(1,43)_ = 10.591 at p=0.002, $: male < female), high-carbohydrate food (center panel; interaction effect of stress and prenatal group, F_(1,43)_ = 5.183 at p=0.028, *: stress PCE > stress CON at p=0.041, #: CON no stress > CON stress), and total combined food (high-fat + high-carbohydrate food; right panel; main effect of stress, F_(1,43)_ = 6.409 at p=0.015, #: no stress > stress; main effect of sex, F_(1,43)_ = 5.567 at p=0.023, $: male < female) during the food choice test, both in no stress and stress conditions. **(C)** 12-hour food choice timepoint intake (kcal/kg) of high-fat food (left panel; main effect of stress, F_(1,43)_ = 10.840 at p=0.002, $: male < female), high-carbohydrate food (center panel; main effect of stress, F_(1,43)_ = 26.424 at p<0.001, #: no stress > stress), and total combined food (high-fat + high-carbohydrate food intake; right panel; interaction effect of sex, stress and prenatal group, F_(1,43)_ = 4.582 at p=0.038, *: male stress PCE > male stress CON at p=0.013, $: CON male < CON female at p=0.007, #: no stress > stress at p<0.01) during the food choice test, both in no stress and stress conditions.

At the 4-hour food choice timepoint, only PCE animals under stress conditions consumed more high-carbohydrate food than control animals (p=0.041, interaction effect of stress and prenatal group, F_(1,43)_=5.183 at p=0.028; Figure 2B). This was not seen in PCE animals under no stress conditions (Figure 2B). Again, no effects of prenatal group were found for high-fat food intake or total combined food intake (Figure 2B).

At the 12-hour food choice timepoint, no effects of prenatal group were found for high-carbohydrate or high-fat food intake in both no stress and stress conditions (Figure 2C). But male PCE animals under stress conditions had higher total combined food intake than male control animals (p=0.013, interaction effect of sex, stress and prenatal group, F_(1,43)_=4.582 at p=0.038; Figure 2C).

As expected, stress reduced total food intake at all timepoints (2-hr: main effect of stress, F_(1,43)_=91.390 at p<0.001; 4-hr: main effect of stress, F_(1,43)_=6.409 at p=0.015; 12-hr: p<0.01, interaction effect of sex, stress and prenatal group, F_(1,43)_=4.582 at p=0.038; Figure 2A-C), and reduced high-carbohydrate food intake (2-hr: main effect of stress, F_(1,43)_=72.820 at p<0.001; 4-hr: p<0.01, interaction effect of stress and prenatal group, F_(1,43)_=5.183 at p=0.028; 12-hr: main effect of stress, F_(1,43)_=26.424 at p<0.001; Figure 2A-C). Females consumed more high-fat food than males at all timepoints (2-hr: p=0.003; interaction effect of sex and stress, F_(1,43)_=4.534 at p=0.039; 4-hr: main effect of sex, F_(1,43)_=10.591 at p=0.002; 12-hr: main effect of sex, F_(1,43)_=10.840 at p=0.002; Figure 2A-C). No significant effects of prenatal group were found for light-phase or 24-hr energy intake (Supplementary Figure 1B-C).

### HFD Access: Weight Gain and Energy Intake

During HFD/LFD diet access, PCE animals gained less weight than control animals from day 12 onwards (p<0.05, interaction effect of day and prenatal group, F_(1.534,59.835)_=4.781 at p=0.019, Figure 3A). All animals gained weight over time with those on the HFD gaining more weight than LFD animals from day 3 onwards (p<0.001, interaction effect of day and diet, F_(1.534,59.835)_=15.160 at p<0.001; Figure 3A). Further, males gained more weight than females at every timepoint (p<0.001, interaction effect of day and sex, F_(1.534,59.835)_=41.118 at p<0.001; Figure 3A).

**Figure 3:**
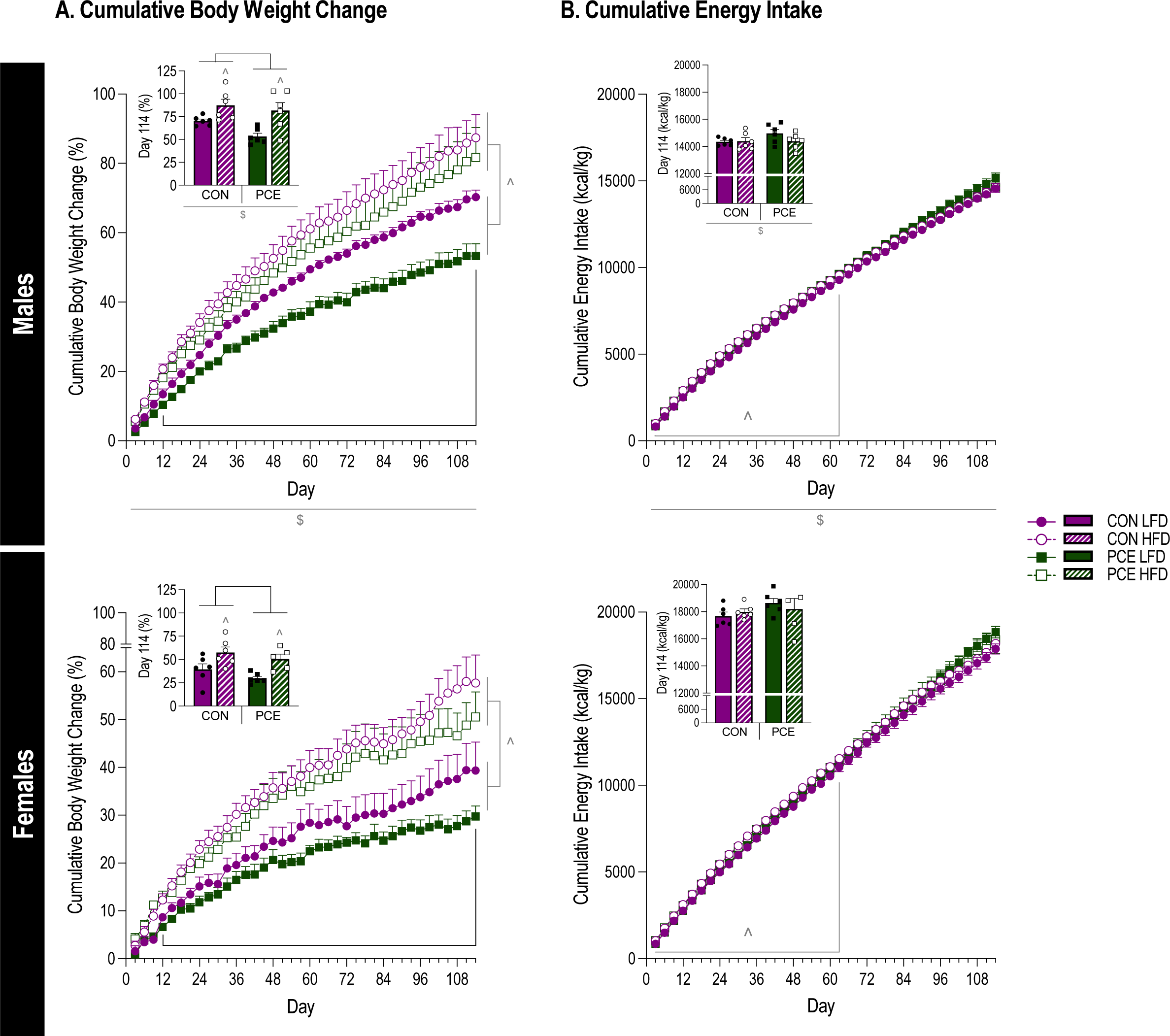
With HFD or LFD access, PCE reduces bodyweight gain over time irrespective of diet but has no effect on energy intake. **(A)** Bodyweight change (%) across the special diet access period for males (top panel) and females (bottom panel; interaction effect of day and prenatal group, F_(1.534,59.835)_ = 4.781 at p=0.019, *: PCE < CON at p<0.05; interaction effect of day and diet, F_(1.534,59.835)_ = 15.160 at p<0.001, ^: HFD > LFD at p<0.001; interaction effect of day and sex, F_(1.534,59.835)_ = 41.118 at p<0.001, $: males > females at p<0.001). Insets: Bodyweight change (%) on day 114 of special diet access (main effect of prenatal group, F_(1,39)_ = 6.474 at p=0.015, *: PCE < CON; main effect of diet, F_(1,39)_ = 29.956 at p<0.001, ^: HFD > LFD; main effect of sex, F_(1,39)_ = 55.771 at p<0.001, $: male > female). **(B)** Cumulative energy intake (kcal/kg) across the special diet access period for males (top panel) and females (bottom panel; interaction effect of day and diet, F_(1.251,48.790)_ = 3.728 at p=0.050, ^: HFD > LFD at p<0.025; interaction effect of day and sex, F_(1.251,48.790)_ = 159.980 at p<0.001, $: males < females at p<0.02). Insets: total energy intake during 114-days of special diet access (main effect of sex, F_(1,39)_ = 226.755 at p<0.001, $: male < female).

For cumulative energy intake, no significant effects of prenatal group were found (Figure 3B). HFD animals consumed more energy than LFD animals between day 3 and 63 (p<0.025, interaction effect of day and diet, F_(1.251,48.790)_=3.728 at p=0.050; Figure 3B). Further, males consumed less energy than females at every timepoint (p<0.02, interaction effect of day and sex, F_(1.251,48.790)_=159.980 at p<0.001; Figure 3B).

### HFD-Testing: Basal Circulating Factors

For basal plasma glucose concentrations, PCE males showed higher glucose levels compared to control males (p=0.039), but no effect was observed for females (interaction effect of prenatal group and sex, F_(1,,47)_=4.282 at p=0.045; Table 1D). Male glucose was higher than females (p<0.03; Table 1D). HFD animals showed lower levels of glucose than LFD animals (main effect of diet, F_(1,47)_=6.173 at p=0.017; Table 1D).

For basal plasma insulin concentrations, control HFD animals showed lower insulin levels than control LFD animals (p<0.001), whereas PCE animals do not show decreased insulin with HFD access (interaction effect of prenatal group and diet, F_(1,47)_=4.060 at p=0.05 Table 1D). Males had higher insulin levels than females (p<0.001; interaction effect of sex and diet, F_(1,47)_=6.172 at p=0.017; Table 1D).

For basal plasma leptin concentrations, no significant effects of prenatal group were found. HFD animals had higher leptin levels than LFD animals (main effect of diet, F_(1,47)_=6.463 at p=0.015), and males had higher levels than females (main effect of sex, F_(1,47)_=25.140 at p<0.001; Table 1D).

### HFD-Testing: Glucose Tolerance

For fasting blood glucose levels (T0), no significant effects of prenatal group were found (Figure 4A). However, following dextrose injection, PCE males had significantly higher glucose levels than control males at T15 and T60 (p<0.05) and trending at T120 (p=0.058; interaction of time, sex and prenatal group, F_(2.568, 100.151)_= 3.33 at p=0.029; Figure 4A); whereas, PCE females had significantly lower glucose levels than control females at T120 (p=0.09; Figure 4A). These patterns are highlighted by area under the curve analysis in Figure 4B.

**Figure 4:**
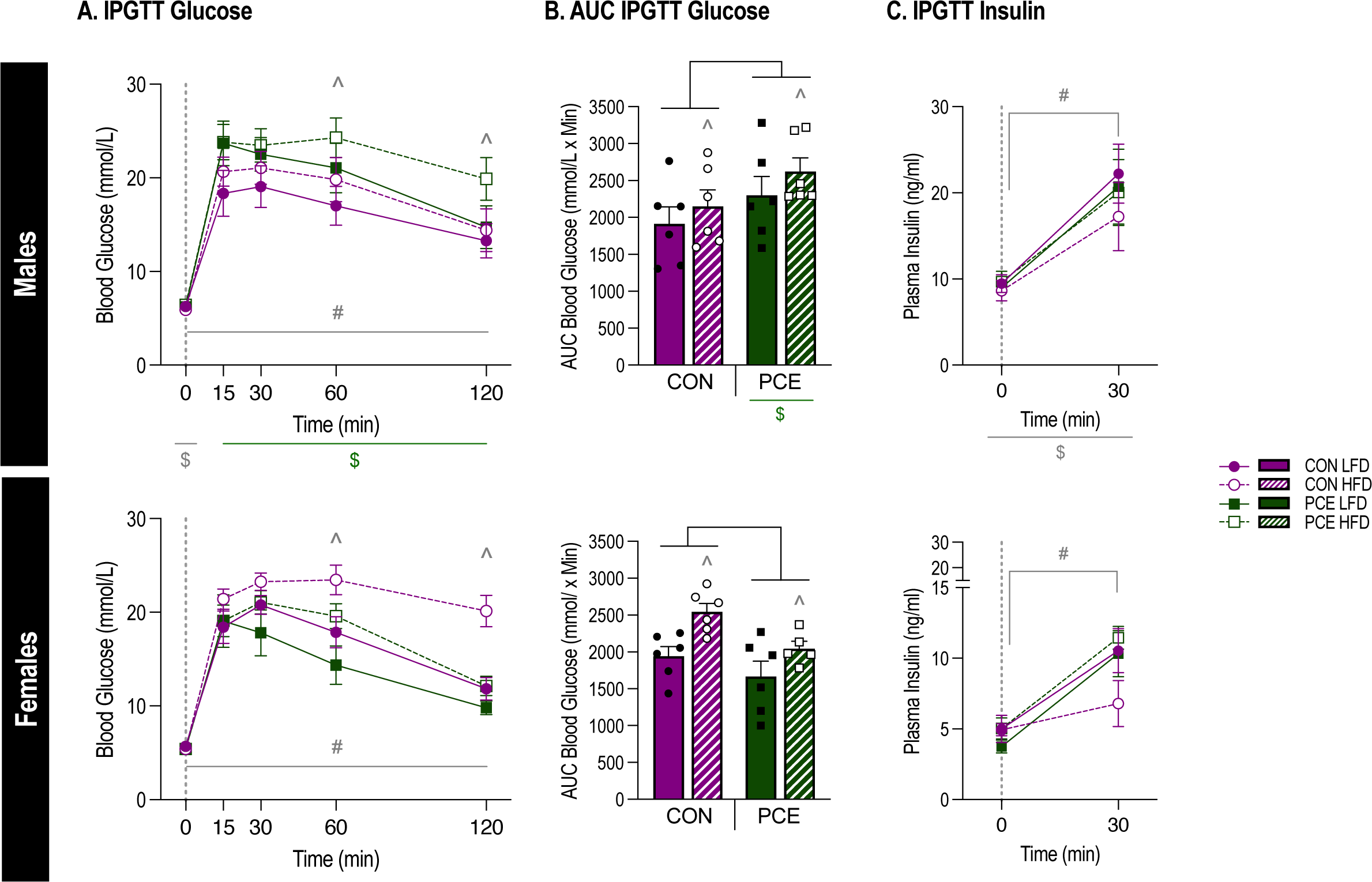
In HFD/LFD-fed offspring, PCE has bi-directional, sex-dependent effects on glucose metabolism, further disrupting glucose metabolism in males and improving glucose metabolism in females. **(A)** Blood glucose (mmol/L) across the IPGTT for males (top panel) and females (bottom panel; interaction of time, sex and prenatal group, F_(2.568, 100.151)_= 3.33 at p=0.029, * (top panel): male PCE > male CON at T15 & T60 at p<0.05, * (bottom panel): female PCE < female CON at T120 at p=0.09, $ (5): CON males > CON females at p=0.004, $ (3): PCE males > PCE females at all time points at p<0.05, #: T0 < T15-T120 at p<0.001; interaction of time and diet, F_(2.568, 100.151)_= 4.506 at p=0.008, ^: HFD > LFD at T60 & T120 at p<0.01). Dotted line indicates timing of dextrose injection. **(B)** AUC IPGTT blood glucose (mmol/L x min) for males (top panel) and females (bottom panel; interaction effect of sex and prenatal group, F_(1, 39)_= 9.155 at p=0.04, * (top panel): male PCE > male CON at p=0.029, * (bottom panel): female PCE < female CON at p=0.05 $: male PCE > female PCE at p=0.003; main effect of diet, F_(1, 39)_= 8.159 at p=0.007, ^: HFD > LFD). **(C)** Plasma insulin (ng/ml) at IPGTT T0 and T30 for males (top panel) and females (bottom panel) (interaction effect of time and sex F_(1, 39)_= 9.966 at p=0.003, #: T30 > T0 at p<0.001, $: male > female at p<0.001). Dotted line indicates timing of dextrose injection.

In general, all animals followed the expected IPGTT time course where glucose increased following dextrose injection (p<0.001) and decreased over time (p<0.01; Figure 4A). Diet did not influence fasting glucose, but following dextrose injection, HFD animals had higher T60 and T120 glucose levels than LFD animals (p<0.01, interaction of time and diet, F_(2.568, 100.151)_=4.506 at p=0.008; Figure 4A). Control males had higher glucose levels than females at T0 (p=0.004), whereas, PCE males had higher glucose levels than females at all timepoints (p<0.05; Figure 4A).

For IPGTT plasma insulin levels, no significant effects of prenatal group or diet were found (Figure 4C). As expected, insulin concentrations were higher after dextrose injection (p<0.001, interaction effect of time and sex F_(1, 39)_=9.966 at p=0.003), and male levels were higher than females (p<0.001; Figure 4C).

For HOMA-IR, there was no significant effect of prenatal group or diet, but males had higher values than females (main effect of sex, F_(1, 39)_=56.126 at p<0.001; data not shown).

### HFD-Testing: Feeding Patterns

Dark-phase energy intake patterns were unaffected by prenatal group or diet, where all rats steadily consumed energy throughout the dark-phase (p<0.02, interaction effect of time and sex, F_(1, 39)_=4.984 at p=0.03; Supplementary Figure 2A). Light-phase and 24-hr energy intake were also unaffected by prenatal group (Supplementary Figure 2B & Figure 5A).

**Figure 5:**
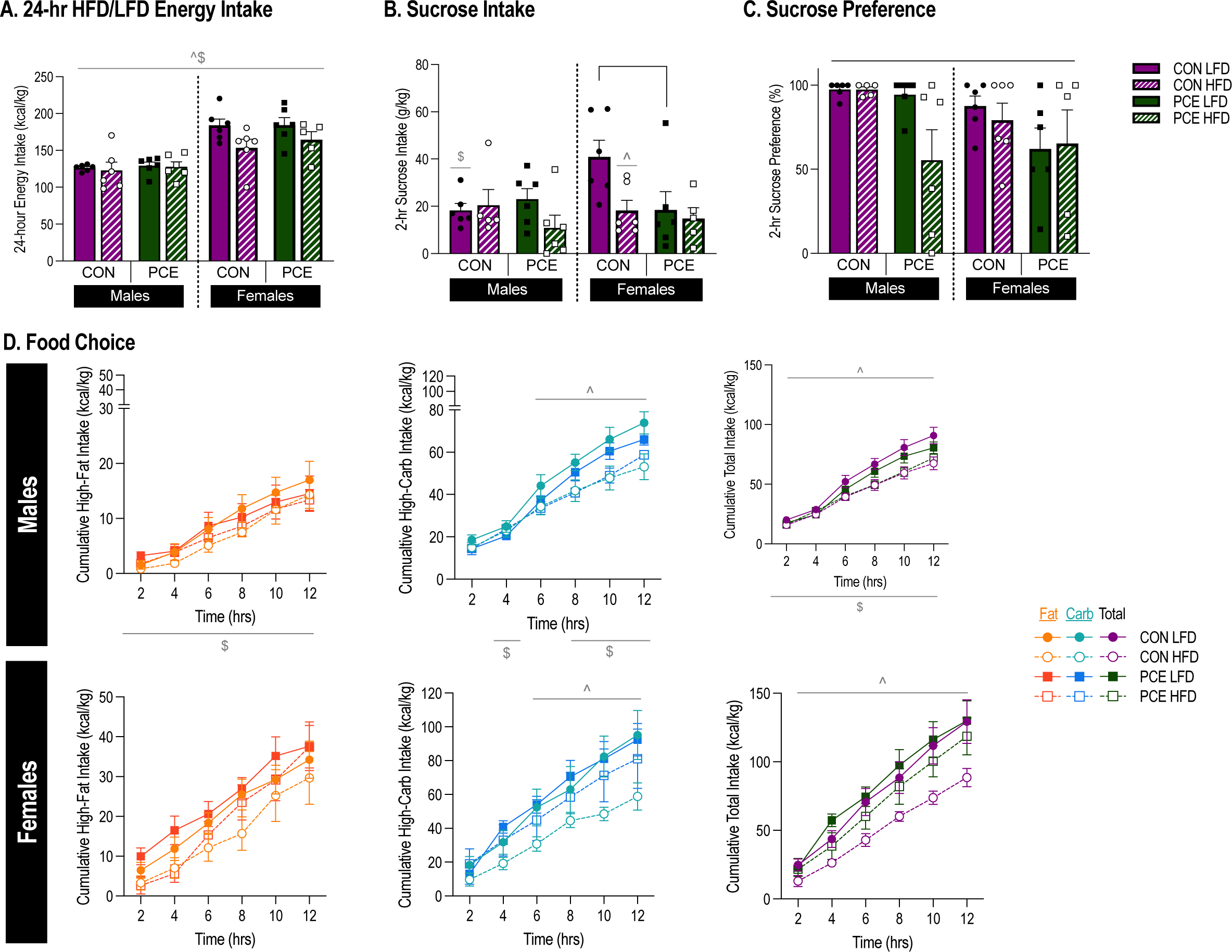
In HFD/LFD-fed offspring, PCE has no effect on energy intake patterns or food choice but reduces sucrose preference. **(A)** *Energy intake patterns:* total 24-hr energy intake (kcal/kg) for HFD- or LFD- fed males and females (main effect of sex, F_(1, 39)_= 53.215 at p<0.001, $: males < females; main effect of diet, F_(1, 39)_= 4.984 at p=0.031, ^: HFD < LFD). **(B)** *Sucrose preference:* 2-hour dark-phase sucrose intake (g/kg; interaction effect of sex, prenatal group, and diet, F_(1, 38)_= 4.438 at p=0.042, *: female LFD PCE < female LFD CON at p=0.006, ^: female CON LFD > female CON HFD at p=0.006, $: male CON LFD < female CON LFD at p=0.006) for males and females. **(C)** *Sucrose preference*: 2-hour dark-phase sucrose preference (%; main effect of prenatal group, F_(1, 38)_= 7.041 at p=0.012, *: PCE < CON). **(D)** *Food choice:* cumulative dark-phase intake (kcal/kg) of high-fat food (left panel; interaction effect of time and sex, F_(1.571,59.681)_= 20.414 at p<0.001, $: male < female at all time points at p<0.001), high-carbohydrate food (center panel; interaction effect of time and sex, F_(2.52,95.745)_= 5.621 at p=0.002, $: male < female at T4, 8, 10 & 12 at p<0.05; interaction effect of time and diet, F_(2.52,95.745)_= 6.188 at p=0.001, ^: LFD > HFD at T6-T12 at p<0.03), and total food (high-fat + high-carbohydrate food; right panel; main effect of diet, F_(1,38)_= 8.786 at p=0.005, ^: LFD > HFD; main effect of sex, F_(1,38)_= 30.077 at p<0.001, $: male < female) over time for males and females during the food choice test.

For dark-phase, light-phase and 24-hr intake, males consumed less food than females (dark-phase: main effect of sex, F_(1, 39)_= 10.784 at p=0.002; light-phase: main effect of sex, F_(1, 39)_=13.892 at p<0.001, 24-hr: main effect of sex, F_(1, 39)_=53.215 at p<0.001; Supplementary Figure 2A-B & Figure 5A), and HFD animals had lower 24-hr intake than LFD animals (main effect of diet, F_(1, 39)_=4.984 at p=0.031; Figure 5A).

### HFD-Testing: Food Choice Test

During the food choice test, there was no significant effect of prenatal group on high-fat, high-carbohydrate, or total combined food intake (high-carbohydrate + high-fat food intake) at any time point (Figure 5D). There was also no effect of prenatal group on light-phase intake (Supplementary Figure 2C). However, for 24-hr energy intake, control LFD animals had higher intake than PCE LFD animals (p=0.022, interaction effect of prenatal group and diet, F_(1, 38)_=4.762 at p=0.035; Supplementary Figure 2D).

HFD animals had lower high-carbohydrate and total combined food intake than LFD animals (high-carbohydrate: p<0.03, interaction effect of time and diet, F_(2.52,95.745)_=6.188 at p=0.001; total intake: main effect of diet, F_(1,38)_=8.786 at p=0.005; Figure 5D). Further, control HFD animals had lower 24-hr energy intake than control LFD animals (p=0.010), an effect not seen in PCE animals (Supplementary Figure 2D). Finally, for all foods and measurements, intake was lower for males than females (high-fat: p<0.001, interaction effect of time and sex, F_(1.571,59.681)_=20.414 at p<0.001; high-carbohydrate: p<0.05, interaction effect of time and sex, F_(2.52,95.745)_=5.621 at p=0.002; total combined intake: main effect of sex, F_(1,38)_=30.077 at p<0.001; light-phase HFD/LFD intake: main effect of sex, F_(1, 38)_= 20.429 at p<0.001; 24-hr intake: main effect of sex, F_(1, 38)_=71.021 at p<0.001; Figure 5D & Supplementary Figure 2C-D).

### HFD-Testing: Sucrose Preference

For sucrose intake, PCE LFD females drink less sucrose than control LFD females (p=0.006; interaction effect of sex, prenatal group, and diet, F_(1, 38)_=4.438 at p=0.042; Figure 5B). In females, control HFD animals drink less sucrose than control LFD animals (p=0.006). Finally, male control LFD animals drink less sucrose than female control LFD animals (p=0.006; Figure 5B).

For sucrose preference, PCE animals have a lower sucrose preference than control animals (main effect of prenatal group, F_(1, 38)_=7.041 at p=0.012; Figure 5C).

### HFD Testing: White Adipose Tissue (WAT) and Digestive Organ Weights

For adiposity, there was no significant effect of prenatal group on any WAT pad mass (Supplementary Table 1) or on adipose to bodyweight ratio (data not shown). As expected, animals on HFD had significantly heavier WAT pads than LFD animals (subcutaneous: main effect of diet, F_(1, 39)_= 14.635 at p<0.001; retroperitoneal: main effect of diet, F_(1, 39)_= 12.914 at p<0.001; gonadal: main effect of diet, F_(1, 39)_=10.185 at p=0.003; total WAT mass: main effect of diet, F_(1, 39)_=15.557 at p<0.001; Supplementary Table 1). HFD animals also have a higher adipose to bodyweight ratio (main effect of diet, F_(1, 39)_=26.071 at p<0.001; data not shown). Further, males had heavier subcutaneous, retroperitoneal, and total WAT mass than females (subcutaneous: main effect of sex, F_(1, 39)_=54.722 at p<0.001; retroperitoneal: main effect of sex, F_(1, 39)_=48.193 at p<0.001; total WAT mass: main effect of sex, F_(1, 39)_=47.361 at p<0.001; Supplementary Table 1). Males also had a higher adipose to bodyweight ratio than females (main effect of sex, F_(1, 39)_=14.790 at p<0.001; data not shown).

## Discussion

Emerging evidence implies that PCE is associated with long-term energy homeostasis and metabolic effects (40, 47), although the evidence surrounding this area, and the directional nature of these effects, is equivocal. The overarching aim of this study was to use a cannabis vapor model to investigate if PCE is associated with increased risk for developing obesity, diabetes and/or abnormal eating habits in adulthood, with additional focus on diet and sex. To our knowledge, this is the first study to assess the energy balance consequences of PCE using a translational vapor exposure model, and the first to explore if PCE alters eating habits, including food choices, in combination with HFD access. In this study, PCE was not associated with changes in bodyweight or adiposity but did decrease the rate of bodyweight gain with special diet (HFD/LFD) access. PCE had prominent bidirectional effects on glucose metabolism that were both diet and sex dependent; when chow-fed, PCE improved glucose metabolism in both sexes, but when special diet-fed, PCE improved glucose metabolism in females irrespective of diet, but impaired glucose metabolism in males irrespective of diet. Moreover, PCE had diet-dependent effects on feeding behaviors: when chow-fed, PCE increased total daily energy intake and acutely increased high-carbohydrate intake in the food choice test, an effect that was prolonged following stress; but when special diet-fed, PCE had no effect on total daily energy intake or high-carbohydrate food intake, but did decrease sucrose preference.

### Birthweight

It is a well-documented clinical observation that PCE is associated with lower birth weight (11, 14, 15), a potential consequence of THC altering placental development and function (20). However, with our vapor PCE model (10% THC, 15min/day, G1-21) we did not observe a reduction in birth weight (at P1), but PCE was associated with reduced bodyweight at P22 regardless of sex. Other rodent models showing PCE-induced fetal growth restriction at P1 typically utilize injection administration (19, 20, 48) and no longer see bodyweight changes by P21 (19, 20). We have previously demonstrated the differences in pharmacokinetic profiles between THC administered via injection compared to vapor inhalation in both non-pregnant and pregnant rats (9, 33). Thereby differences in route of administration, pharmacokinetics and/or dose may explain inconsistencies in the effects of PCE on birth weight between rodent models. In fact, consistent with our findings, Benevenuto *et al* 2017 also did not observe overall litter weight changes with a mouse whole cannabis smoke inhalation model (0.3% THC, 5min/day, G5.5-17.5), and Weimar *et al.* 2020 also did not observe THC-specific bodyweight changes with a rat whole cannabis vapor model (24.8% THC, 60min/twice daily, G0-22). Furthermore, several clinical studies have also failed to find an association between PCE and lower birth weight (49). Aligning with other injection and inhalation models, we did not observe any effects of PCE on maternal bodyweight or food intake (17, 20) indicating that maternal nutrition was not a contributing factor.

### Adiposity

Clinical studies imply that PCE is associated with higher BMI and fat mass during childhood (14, 15, 39) and this abnormal ‘catch-up growth’ is hypothesized to increase the risk for developing obesity later in life (47). In our chow-fed rats, PCE had no effect on bodyweight or weight gain during late adolescence (P50) and early adulthood (P77), as well as no effect on plasma leptin (indirect measurement of adiposity). Previous studies have similarly shown no effect of PCE on bodyweight later in life (19) but other studies have shown that PCE increases visceral adipose tissue mass (41). However, due to the longitudinal nature of our study we were unable to directly measure total fat mass when chow-fed. Our indirect measurements imply that our chow-fed PCE animals did not show adiposity changes.

Rats were then placed on an obesogenic HFD (or LFD, starting at P78) to see if the effects of PCE on adiposity would become more apparent. As expected, four-months of HFD increased bodyweight, plasma leptin levels, WAT mass, and impaired glucose metabolism, consistent with other studies (50, 51). During the 4-month “HFD access” period, PCE reduced the rate of bodyweight gain such that by 6-months of age PCE animals had gained less weight than controls (irrespective of diet or sex). PCE did not influence plasma leptin concentrations, WAT mass, or adiposity to bodyweight ratio, suggesting that these bodyweight gain differences were not due to changes in adiposity. Thereby, in our study PCE did not appear to amplify the obesogenic effects of long-term HFD access in adulthood.

Interestingly, THC reducing bodyweight gain has been observed in other rodent models of cannabis exposure and HFD access during adolescence/adulthood (52, 53). Lin *et al* 2023 showed that adolescence THC exposure is associated with reduced bodyweight gain, decreased fat mass, increased lean mass and increased energy expenditure with HFD access in adulthood (53). It is not known if the PCE-induced decrease in bodyweight gain observed in the present study is due to changes in lean body mass or energy expenditure as we did not take these measures.

### Diabetes

There is clinical evidence that blood glucose levels are elevated in childhood following PCE (14), and PCE-induced alterations in glucose metabolism have been reported in rodent studies (19, 54), suggesting that PCE may increase risk for diabetes development later in life. In line with other studies, we showed that in chow-fed young adult rats (P77), PCE did not influence basal or fasting blood glucose or plasma insulin concentrations in either sex (19, 54). However, using an IPGTT, we found that PCE was associated with a subtle improvement of glucose metabolism in both sexes when chow-fed, where peak blood glucose levels were reduced with PCE, but the overall time course and general rate of glucose clearance was unaffected. In contrast, using a THC injection PCE model, Gilles *et al* 2020 showed that PCE impaired glucose metabolism specifically in chow-fed older adult females (19). Further, they showed that in females, increased blood glucose during the IPGTT was accompanied by an increase in peak plasma insulin concentrations, decreased insulin sensitivity, decreased fasting serum glucagon concentrations and changes in pancreatic morphology (19, 54). Together, our studies show that PCE has marked effects on glucose metabolism in chow-fed adults, however, the discrepancies between findings are likely due to differences in route of administration, dose, or testing age.

Interestingly, when we put our animals on HFD or LFD for 4-months, the effects of PCE on glucose metabolism changed in a sex and diet dependent manner. As expected, HFD generally increased the time course of glucose clearance during the IPGTT, demonstrating impaired glucose metabolism, consistent with previous studies using the same diet (50, 55). In males, irrespective of diet, PCE increased peak glucose levels and prolonged glucose clearance, implying that PCE impairs glucose metabolism in males. In females, the opposite was observed, where irrespective of diet, PCE had no effect on peak glucose levels, and accelerated glucose clearance, implying that PCE improves glucose metabolism in females. This female PCE effect did not appear to be influenced by diet. Sex specific effects on glucose metabolism have been reported with other PCE rodent models (19, 54), demonstrating how robust this sex-divergence is.

In addition, basal blood glucose was significantly elevated in male PCE animals irrespective of diet, indicating glucose dysregulation. Further, HFD reduced basal insulin levels irrespective of sex, suggesting that HFD access interfered with insulin production or secretion. Interestingly, this reduction in insulin with HFD was not observed in PCE animals, suggesting that PCE may be having protective effects against HFD in basal conditions. However, there was no difference in plasma insulin concentrations with diet and PCE during the IPGTT, suggesting that PCE effects on glucose metabolism observed during the IPGTT are not caused by changes in insulin secretion. Instead, PCE could be altering insulin sensitivity; Gilles *et al* 2020 showed that PCE alters insulin sensitivity and islet cell morphology in adulthood, specifically reducing the mass of insulin-secreting β-cells in females (19). Further, CB1R are expressed on both insulin-secreting β-cells and glucagon-secreting α-cells within the pancreas (56) and are involved in the survival and organization of pancreatic endocrine islet cells (57, 58) as well as insulin receptor signaling (59). Thus, it is possible that THC could act on the developing pancreas through CB1R leading to changes in glucose metabolism later in life.

### Eating Habits

Many prenatal factors, for example prenatal stress and maternal nutrition, alter feeding behaviors later in life (60, 61). As PCE is potentially associated with increased adiposity and risk for obesity later in life and the underlying mechanism for this remains unknown, it is possible that PCE could disrupt energy balance through promoting abnormal eating patterns. In line with this idea, there is evidence that PCE alters food reward-related behaviors, and this is associated with epigenetic changes in the striatal nucleus accumbens, which is involved in controlling motivated behaviors, including eating (42). Therefore, we investigated if PCE influenced daily energy intake patterns and food choice in adulthood.

### Daily Energy Intake Patterns

Previous studies using injection have shown that PCE has no effect on daily chow intake (41), or on total daily intake of chocolate pellets acquired during a fixed ratio operant task (42) in late adolescent rats (P50–60 and P62 respectively). However, with our vapor PCE model, we found that PCE was associated with increased 24-hr energy intake, specifically through increased light-phase intake, in early adult (P77) chow-fed rats. Thereby, PCE altered daily chow intake patterns. Interestingly, the increase in daily energy intake was not accompanied by increased bodyweight or plasma leptin levels, suggesting that compensatory energy balance mechanisms, for example increased energy expenditure, accounted for higher energy intake. However, we did not measure locomotion, thermogenesis, respiration, or other measures of energy expenditure in this study. It is not known if PCE alters energy expenditure in adulthood, but there is evidence that cannabis exposure during adolescence is associated with increased thermogenesis in adulthood (53). Differences in findings between the present study and previously published studies could be due to route of administration, dose, or testing age. Unlike when chow-fed, when on the special diets (HFD or LFD), PCE was not associated with changes in daily energy intake patterns, thereby the effect of PCE on daily energy intake patterns could be dependent on many variables including diet and age.

### Food Choice

When chow-fed adults were given a choice between a high-carbohydrate food and a high-fat food, PCE rats consumed more energy overall in the first 2-hrs of the food choice test compared to controls and this was mainly through increased high-carbohydrate food intake. However, PCE did not affect the total amount of high-carbohydrate and high-fat food eaten during the entire test or total light-phase and 24-hr energy intake. Thereby, in these conditions, PCE acutely influences intake patterns of palatable, macronutrient-specific foods. It has previously been shown that PCE significantly increased breakpoint to acquire chocolate sugar pellets in a progressive ratio operant task (42), suggesting that PCE is associated with heightened motivation to obtain palatable, high-carbohydrate foods. The acute increases in high-carbohydrate food intake seen in our study could reflect PCE rats having increased motivation for carbohydrates, however, more behavioral testing is required to directly assess this.

Unlike when chow-fed, when on the special diets (HFD/LFD), PCE was not associated with changes in high-fat and high carbohydrate food intake in the food choice test. However, when subjected to an acute 2-bottle choice test to specifically assess sucrose preference, PCE reduced sucrose preference irrespective of sex or diet. Decreased sucrose preference in a similar 2-bottle choice test was previously observed with PCE in animals previously exposed to a stressor (42). Male PCE animals show multiple signs of glucose dysregulation during the ‘HFD testing’ phase of the experiment, therefore decreased sucrose preference in males could be a result of this. However, this does not explain why females also show reduced sucrose preference. More research is required to assess the interaction between PCE, diet and carbohydrate preference, with focus on metabolic and reward mechanisms.

### Environmental Challenges (Stress and HFD)

The ‘double hit hypothesis’ implies that following a prenatal insult (such as PCE), developmental changes occur in many systems in anticipation of future challenges. Thereby, these systems are primed to respond differently to additional insults later in life (62). We assessed whether PCE has a priming effect on energy balance and metabolism after exposure to secondary insults (acute stress and obesogenic HFD).

#### Restraint Stress

Using an intravenous PCE model, it has been shown that PCE rats previously subjected to a forced swim test, i.e., a stressful experience, had reduced sucrose preference compared to rats not subjected to a forced swim test (42), suggesting an interaction between PCE and stress which alters carbohydrate consumption. Therefore, we assessed if stress altered the effect of PCE on food choice, by subjecting chow-fed rats to a 30-min restraint stress and giving them a choice between high-carbohydrate and high-fat food. Restraint stress decreased overall food intake as expected (63, 64) and prolonged the effects of PCE on food choice, where stressed PCE rats ate more high-carbohydrate food intake within the first 4-hrs of the test, compared to no stress conditions where intake was increased in the first 2-hrs only. Stressed PCE male rats also ate more food overall in the 12-hr food choice test, which was not observed in no stress conditions. Thereby an interaction between PCE and stress exists, where acute stress prolongs the effects of PCE on food choice.

#### High-Fat Diet

To our knowledge, no study to date has investigated the consequences of PCE with HFD in adulthood. As expected, HFD access was associated with increased bodyweight gain, increased plasma leptin concentrations (50, 51) and increased time course for glucose tolerance (50, 55) compared to animals on LFD. Throughout the HFD-testing phase of the study, interactions between PCE and HFD access became apparent. For example, the impairing effects of PCE on glucose tolerance in males were moderately amplified by HFD, suggesting an additive effect between the two conditions, where HFD could be exacerbating the potential pancreatic effects of PCE. Further, reductions in basal insulin levels associated with HFD were abolished in PCE animals, implying that the ability of HFD to impair insulin production and/or secretion, was somewhat protected in animals exposed to PCE. Finally, decreased sucrose consumption in female PCE animals on LFD was not seen in female PCE animals on HFD, suggesting that HFD could be disrupting the potential reward-related alterations of PCE. From previous literature, it appears that PCE can induce glucose dysregulation through similar mechanisms to HFD (19), thereby it is not entirely surprising that interactions between PCE and HFD are seen with respect to glucose metabolism.

### Conclusions

Clinical PCE studies are experimentally limited, therefore animal research is critical for assessing the potential long-term energy balance consequences of PCE. This work will enhance current understanding of the potential risks of PCE, knowledge that may be used to develop more accurate public health guidelines to better advise individuals on the use of cannabis during pregnancy.

## Acknowledgements

We would like to acknowledge that Indigenous peoples are the original and current caretakers of the land we live, work, learn, and play on. Specifically, our research was conducted on the traditional territories of the people of the Treaty 7 region in Southern Alberta, which includes the Blackfoot Confederacy (comprising the Siksika, Piikani, and Kainai First Nations), as well as the Tsuut’ina First Nation, and the Stoney Nakoda (including the Chiniki, Bearspaw, and Wesley First Nations). The City of Calgary is also home to Métis Nation of Alberta, Region III. As an academic community, we must recognize that their land was taken through coercive and violent acts, and we must support their authority and rights over this stolen land. We must acknowledge our responsibility to establish and maintain relationships with Indigenous peoples and we must include their voices in our teaching and research.

We would also like to thank Lauren Seabrook for advice and assistance with data analysis and glucose tolerance testing; Georgia Balsevich for experimental design advice; Andrei (Sabin) Nastase for help with data collection; and finally, Robert Aukema and Gavin Petrie for help with the glucose tolerance testing.

