## Supplementary Figures for "Prenatal exposure to THC vapor influences feeding, bodyweight and glucose metabolism under both basal conditions and following high fat diet"

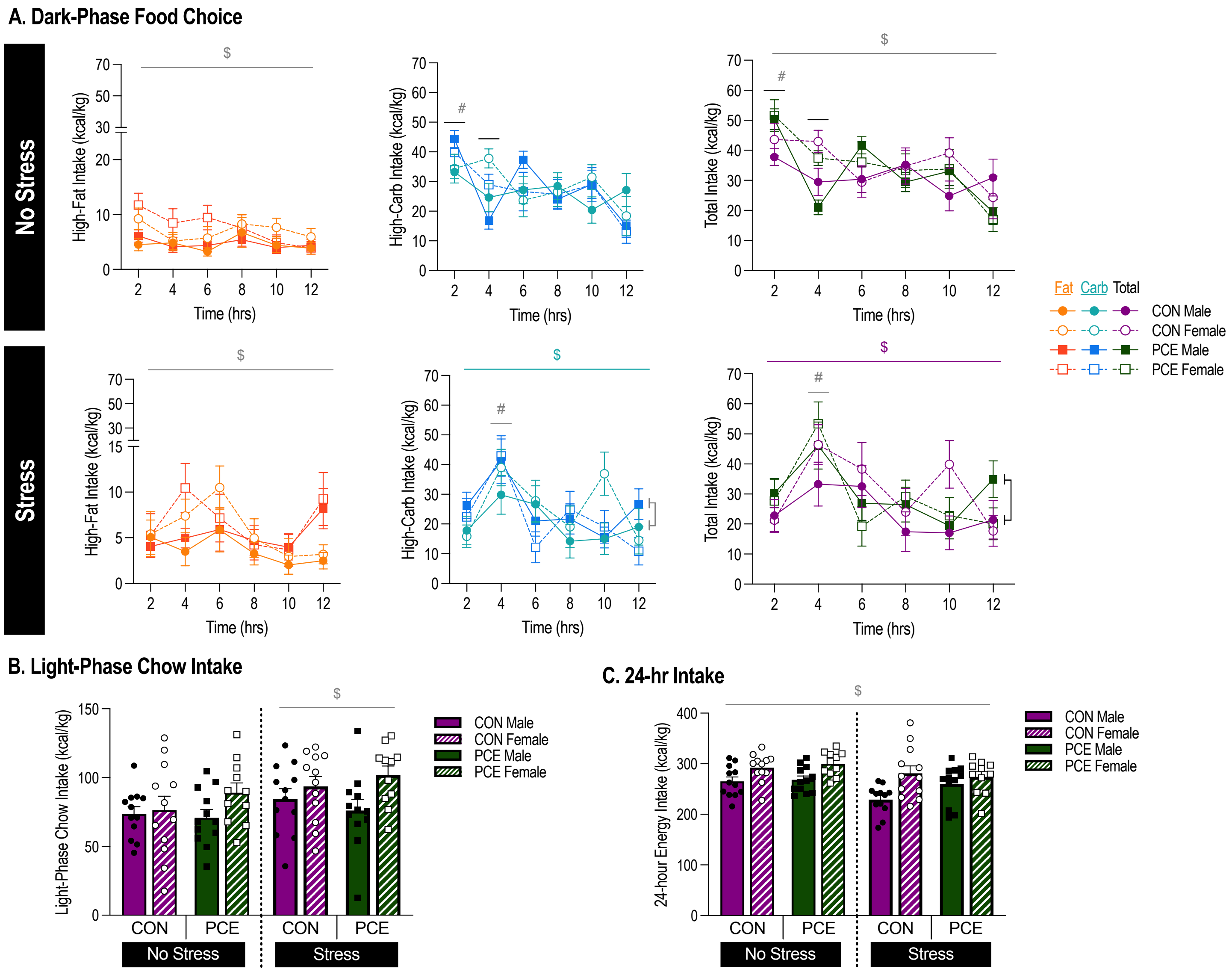


**Supplementary Figure 1: In chow-fed offspring, PCE and stress acutely alters intake patterns of high-carbohydrate food in a 12-hour dark-phase food choice test but does not alter light-phase chow intake or 24-hr energy intake during this test.**

**(A)** Top panel: dark-phase food choice test in *no stress* conditions: intake (kcal/kg) of high-fat food (left panel; main effect of sex, $F_{(1,43)} = 14.783$ at p<0.001, $: male < female), high-carbohydrate food (center panel; interaction effect of time & prenatal group, $F_{(3.708,159.441)} = 2.483$ at p=0.05, *: T2 PCE > T2 CON at p=0.035, *: T4 PCE < T4 CON at p=0.029; main effect of time, $F_{(3.708,159.441)} = 7.126$ at p<0.001, #: T2 > T4, 8, 10 & 12 at p<0.05), and total combined food (high-fat + high-carbohydrate food; right panel; interaction effect of time & prenatal group, $F_{(3.849,165.526)} = 2.734$ at p=0.033, *: T2 PCE > T2 CON at p=0.023, *: T4 PCE < T4 CON at p=0.049; main effect of time, $F_{(3.849,165.526)} = 8.544$ at p<0.001, #: T2 > all other time points at p<0.03; main effect of sex, $F_{(1,43)} = 4.249$ at p=0.045, $: male < female) over time during the food choice test. Bottom panel: dark-phase food choice test in *stress* conditions: intake (kcal/kg) of high-fat food (left panel; main effect of sex, $F_{(1,43)} = 4.318$ at p=0.044, $: male < female), high-carbohydrate food (center panel; interaction effect of sex & prenatal group, $F_{(1,43)} = 6.020$ at p=0.018, *: male PCE > male CON at p=0.045, $: CON male < CON female at p=0.039; main effect of time, $F_{(4.099,176.269)} = 6.102$ at p<0.001, #: T4 > all other time points at p<0.05), and total combined food (high-fat + high-carbohydrate food; right panel; interaction effect of sex & prenatal group, $F_{(1,43)} = 6.288$ at p=0.016, *: male PCE > male CON at p=0.013, $: CON male < CON female at p=0.007; main effect of time, $F_{(5,215)} = 5.762$ at $p<0.001$, #: T4 > T2, 8, 10 & 12 at $p<0.05$) over time during the food choice test following restraint stress. **(B)** Total 12-hour light-phase chow intake (kcal/kg) during the food choice test in *no stress* conditions (no statistically significant effects) and following restraint *stress* (main effect of sex, $F_{(1,43)} = 5.601$ at $p=0.023$, $: male < female). **(C)** 24-hr intake (kcal/kg; dark-phase high-fat & high-carbohydrate food intake + light-phase chow intake) during the food choice test in *no stress* conditions (main effect of sex, $F_{(1,43)} = 14.207$ at $p<0.001$, $: male < female) and following restraint *stress* (main effect of sex, $F_{(1,43)} = 8.462$ at $p=0.006$, $: male < female).

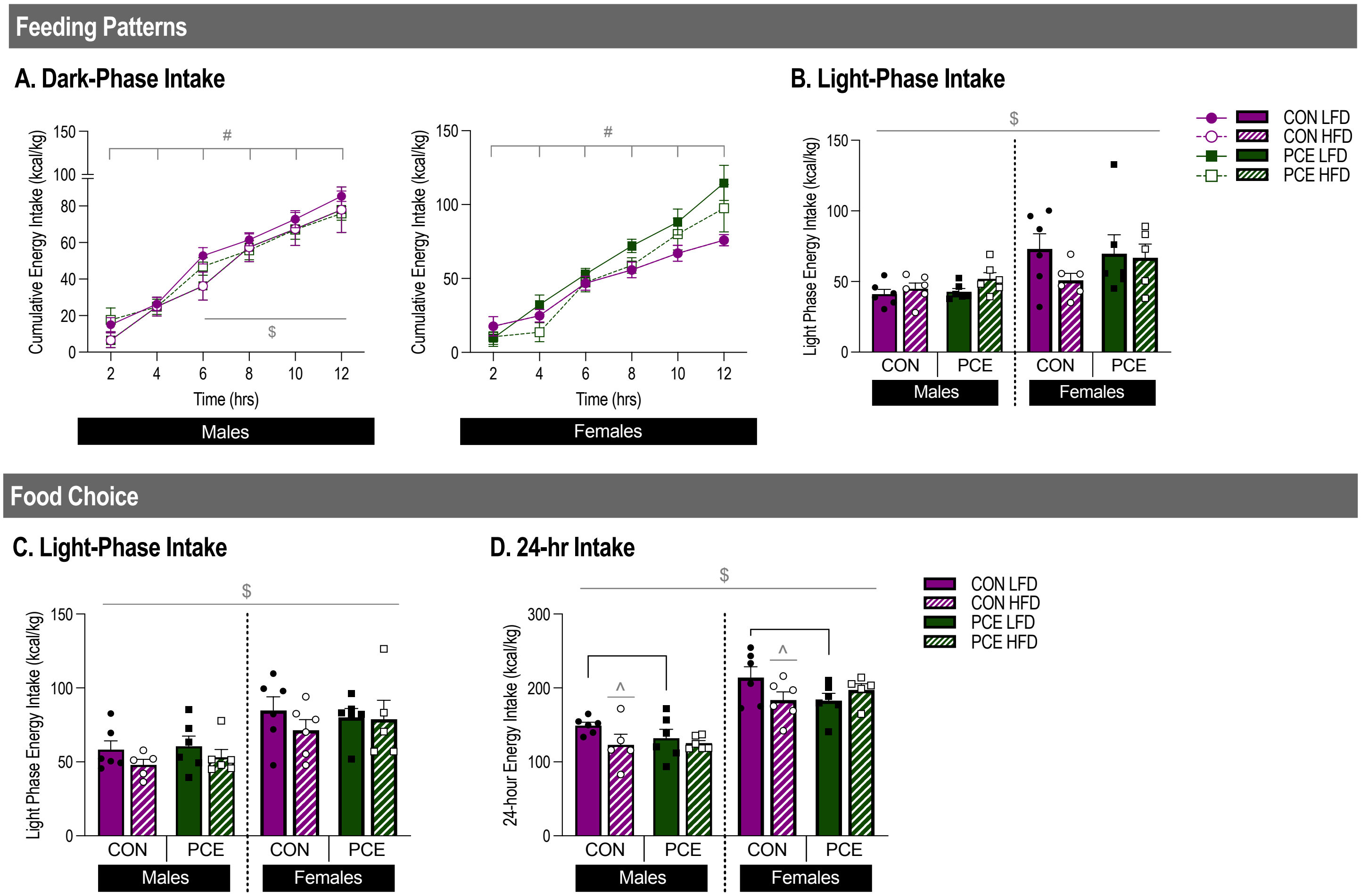


**Supplementary Figure 2: In HFD/LFD-fed offspring, PCE has no effect on dark or light-phase intake patters but does decrease 24-hr energy intake in LFD-fed animals during a food choice test.** *HFD/LFD energy intake patterns:* **(A)** Cumulative dark-phase energy intake (kcal/kg) over time for males and females (interaction effect of time and sex, $F_{(1, 39)}$= 4.984 at p=0.03, #: all timepoints significantly different than all other timepoints at p<0.02, $: males < females at T6-T12). **(B)** Total 12-hour light-phase energy intake (kcal/kg) for males and females (main effect of sex, $F_{(1, 39)}$= 13.892 at p<0.001, $: males < females). *Food choice:* **(C)** Total 12-hour light-phase HFD/LFD intake (kcal/kg) for males and females during the food choice test (main effect of sex, $F_{(1, 38)}$= 20.429 at p<0.001, $: male < female). **(D)** 24-hr intake (kcal/kg; dark-phase high-fat & high-carbohydrate food intake + light-phase HFD/LFD intake) for males and females during the food choice test (interaction effect of prenatal group and diet, $F_{(1, 38)}$= 4.762 at p=0.035, *: LFD CON > LFD PCE at p=0.022, ^: CON LFD > CON HFD at p=0.010; main effect of sex, $F_{(1, 38)}$= 71.021 at p<0.001, $: male < female).

| White Adipose Tissue (WAT) Weights | | | | | | | | |
|---|---|---|---|---|---|---|---|---|
| Organ | LFD | | | | HFD | | | |
| | Male | | Female | | Male | | Female | |
| | CON | PCE | CON | PCE | CON | PCE | CON | PCE |
| Subcutaneous WAT (g) $^{\$\wedge}$ | 41.9 ± 30.4 | 38.5 ± 38.0 | 12.4 ± 20.9 | 9.9 ± 24.2 | 68.9 ± 57.8 | 66.9 ± 80.7 | 20.9 ± 25.9 | 23.4 ± 60.4 |
| Retroperitoneal WAT (g) $^{\$\wedge}$ | 37.9 ± 0.3 | 30.1 ± 0.4 | 8.8 ± 0.3 | 10.1 ± 0.3 | 56.1 ± 0.4 | 64.6 ± 0.6 | 18.1 ± 0.5 | 17.5 ± 0.5 |
| Gonadal WAT (g) $^{\wedge}$ | 15.8 ± 4.1 | 15.6 ± 3.6 | 11.3 ± 1.4 | 12.0 ± 0.7 | 20.1 ± 12.3 | 22.1 ± 12.0 | 16.8 ± 3.9 | 19.5 ± 7.9 |
| Total Fat (g) $^{\$\wedge}$ | 95.6 ± 5.2 | 84.2 ± 6.3 | 32.5 ± 0.7 | 32.0 ± 2.0 | 145.0 ± 10.0 | 153.6 ± 12.9 | 55.9 ± 3.0 | 60.4 ± 4.9 |

**Supplementary Table 1: White Adipose Tissue (WAT) Weights**

Table shows weights of subcutaneous WAT (main effect of diet, $F_{(1, 39)}$= 14.635 at p<0.001, ^: HFD > LFD; main effect of sex, $F_{(1, 39)}$= 54.722 at p<0.001, $: male > female), retroperitoneal WAT (main effect of diet, $F_{(1, 39)}$= 12.914 at p<0.001, ^: HFD > LFD; main effect of sex, $F_{(1, 39)}$= 48.193 at p<0.001, $: male > female), gonadal WAT (main effect of diet, $F_{(1, 39)}$= 10.185 at p=0.003, ^: HFD > LFD) and total WAT (main effect of diet, $F_{(1, 39)}$= 15.557 at p<0.001, ^: HFD > LFD; main effect of sex, $F_{(1, 39)}$= 47.361 at p<0.001, $: male > female), for male and female, PCE and control animals fed HFD or LFD.
